# Privacy-Preserving Genomic Data Sharing via Hybrid AES/ECC and Blockchain-Based Dynamic Consent

**DOI:** 10.1101/2025.11.01.686038

**Authors:** Joe Soundararajan, Yan Zhuang, Dong Xu

**Affiliations:** Institute for Data Science and Informatics, University of Missouri, Columbia, MO 65201, USA; Department of Biomedical Engineering and Informatics, Luddy School of Informatics, Computing, and Engineering, Indiana University, Indianapolis, IN 46202, USA; Department of Electrical Engineering and Computer Science, College of Engineering, University of Missouri, Columbia, MO 65201, USA; Bond Life Sciences Center, University of Missouri, Columbia, MO 65201, USA

**Keywords:** Blockchain, Genomics, Ethereum, Hybrid AES/ECC, InterPlanetary File System

## Abstract

Genomic data are growing exponentially, creating opportunities for discovery but also raising significant concerns regarding privacy, security, and governance. We present a framework that encrypts genome-scale files using AES-256-GCM and protects keys per recipient via Curve25519, anchors only content identifiers and encrypted keys on a blockchain smart contract, and stores ciphertext off-chain in IPFS. The design enables dynamic, auditable consent through on-chain access-control lists (grant/revoke with immutable events) while keeping raw genomes off-chain. A working prototype on the Ethereum Sepolia testnet demonstrates end-to-end registration, retrieval, and revocation. Experiments on the human reference genome (hg38) and synthetic sequences demonstrate near-AES performance for multi-gigabyte files and strong diffusion, as per the avalanche criterion. By separating the data plane (symmetric encryption) from the control plane (public-key keying + smart contracts), the system delivers practical, privacy-preserving genomic data sharing suitable for IRB-approved cohort and clinical workflows.

## 1. Introduction

The field of genomics has experienced unprecedented growth in recent years, driven by advancements in high-throughput sequencing technologies. This exponential increase in genomic data generation presents immense opportunities for biomedical research and significant challenges in data management, security, and privacy [1]. The volume of genomic data produced is outpacing Moore’s Law, creating substantial hurdles for storage, processing, and analysis [2]. Current centralized systems struggle to scale efficiently with the increasing volume of data, necessitating novel approaches to genomic data management [3]. One of the primary concerns in genomic data handling is its sensitive nature. Human genomic data is highly personal, revealing information about individuals and their relatives [4]. This raises critical privacy and security concerns, particularly in light of regulations such as the General Data Protection Regulation (GDPR) in the European Union and the Health Insurance Portability and Accountability Act (HIPAA) in the United States [5]. Another significant challenge is striking a balance between data accessibility for research and individual privacy rights. The scientific community recognizes the value of data sharing to advance genomic research and personalized medicine. However, current systems often lack fine-grained, dynamic access control mechanisms that can adapt to the evolving needs of researchers while respecting the privacy preferences of individuals [6]. Ensuring the authenticity and traceability of genomic data throughout its lifecycle is also crucial. Many existing solutions struggle to provide tamper-proof audit trails of data access and modifications, which is essential for maintaining data integrity and provenance [7].

Blockchain technology, originally developed as the underlying system for cryptocurrencies like Bitcoin, has emerged as a promising solution for data management challenges in genomics [8]. At its core, blockchain is a decentralized, distributed ledger that records transactions across a network of computers. This technology offers several key features that make it particularly suitable for genomic data management [9]. The decentralized nature of blockchain means no single entity controls the entire network, reducing the risk of data manipulation or single points of failure [10]. Once data is recorded on the blockchain, it is extremely difficult to alter, ensuring data integrity through immutability [11]. Furthermore, all transactions are visible to network participants, promoting trust and accountability through transparency [12]. Blockchain also enables smart contracts, self-executing contracts with terms directly written into code, facilitating automated and secure data-sharing agreements [13]. In the context of genomic data management, these blockchain features translate into several potential benefits. Every access to genomic data can be recorded immutably on the blockchain, creating a transparent and tamper-proof audit trail [14]. This enhances data ownership, allowing individuals greater control over their genomic data and deciding who can access it and for what purposes [15]. Implementing smart contracts can facilitate secure and automated data sharing, potentially accelerating collaborative genomic research [16]. The immutable nature of blockchain ensures the traceability of genomic data access throughout its lifecycle, maintaining data provenance [17]. Moreover, blockchain can serve as a unified platform for managing genomic data across different healthcare systems and research institutions, improving interoperability in the field [18].

### Current Gaps

While blockchain provides an immutable, decentralized, and transparent framework for securing genomic data transactions, it is not designed to handle large genomic datasets. Whole-genome sequences can be hundreds of gigabytes in size, making direct on-chain storage prohibitively expensive and inefficient due to the blockchain’s inherent limitations in throughput and storage costs. To address this challenge, our approach integrates blockchain for metadata and access control, while leveraging the InterPlanetary File System (IPFS), a peer-to-peer protocol and network designed for decentralized data storage and sharing, particularly for off-chain genomic data storage [19]. This hybrid model ensures that the integrity, traceability, and security benefits of blockchain are maintained while allowing for scalable and cost-effective storage of large genomic datasets. Our blockchain-integrated genomic data management framework combines IPFS for decentralized storage, hybrid encryption for privacy and security, and smart contract-based access control to enforce fine-grained permissions. The framework tokenizes genomic data by storing cryptographic metadata on-chain, such as IPFS hashes (content identifiers) and other essential information that uniquely represent the genomic data without exposing the raw data itself. Genomic tokenization involves replacing sensitive genomic data elements with unique tokens that preserve the data’s utility for research while protecting individual privacy. Tokenization allows researchers to work with genomic data without accessing raw, identifiable information, thereby reducing privacy risks. Furthermore, tokenized data can be more freely shared among researchers and institutions, facilitating collaborative studies while maintaining compliance with privacy regulations [20]. The blockchain records these tokens along with access control policies, creating an immutable audit trail of data usage.

The integration of IPFS with blockchain offers several advantages. First, it enables decentralized genomic data storage, eliminating the need for centralized storage solutions vulnerable to single points of failure and data breaches [21]. Second, IPFS’s content-addressing mechanism ensures that data is uniquely identified based on its content rather than its location, making it resistant to tampering and enabling efficient data retrieval. To address data security, advanced cryptographic techniques are being explored, with hybrid encryption schemes showing promise. The combination of symmetric and asymmetric encryption, specifically the hybrid use of Advanced Encryption Standard (AES) and Elliptic Curve Cryptography (ECC), offers a robust solution for genomic data security [22]. A hybrid encryption approach combining AES (symmetric) and ECC (asymmetric) was deliberately chosen after considering several alternatives. AES-256 was selected for encrypting the bulk genomic data due to its proven efficiency and security for large files. Symmetric ciphers like AES are well-suited for big data because of their high throughput. AES encryption is hardware-accelerated on most modern processors, offering faster performance than older algorithms. For asymmetric encryption and key exchange, we opted for ECC over the more traditional RSA algorithm [23] for efficiency: ECC offers equivalent security with much smaller keys, which is beneficial in a blockchain context where every byte of stored data (and every computation) can cost. Smaller key sizes also translate into faster cryptographic operations (e.g., signature verification, encryption/decryption), which is especially important when a smart contract must handle many access requests. ECC’s computational speed and low resource usage have been noted to generally outperform RSA at common security levels [24]. We also note that this hybrid approach is forward-looking: it can be extended or upgraded if needed, addressing long-term concerns about cryptographic resilience, whereas older ciphers like 3DES or RSA-2048 are more vulnerable to future advances in computing [25].

### Related Works

This paper primarily focuses on evaluating the encryption mechanisms and integration of blockchain technology for secure genomic data management, rather than assessing blockchain’s performance metrics such as throughput or latency. Our contribution is a systems/engineering integration of established components (AES for bulk encryption, ECC for key protection, IPFS for off-chain storage, and Ethereum smart contracts for access/audit) tailored to genomic privacy. We do not introduce new cryptographic primitives; instead, we implement and evaluate an end-to-end workflow for whole-genome–scale sharing with consent-aware access control. By employing blockchain and IPFS together, this framework addresses the pressing challenges of genomic data security, privacy, and scalability, while facilitating transparent and secure data sharing across the biomedical research community [26].

Existing research on secure genomic data management has begun exploring blockchain technology and advanced encryption. Several studies have applied blockchain to general health data and even genomic scenarios. For instance, Dambrot et al. introduced ReGene, a blockchain-based system to back up and share genomic information [27]. Their approach utilizes a distributed ledger to ensure that genomic data and associated metadata can be recovered and verified, highlighting the potential of blockchain for data integrity in genomics. Furthermore, Jin et al. describe a blockchain-powered platform (LifeCODE.ai) for personal genomic data sharing, demonstrating how decentralization, traceability, and encryption can address issues like data tampering and privacy leakage [28]. Their results show that blockchain opens new avenues for genomic data ownership and security, empowering individuals with greater control over how their DNA data is used. Similarly, Park et al. propose a blockchain-based genomic management system that combines on-chain access control with off-chain storage and local differential privacy, adding random noise to genomic data to protect sensitive variants [29]. This approach enables the sharing of genomic datasets in a privacy-preserving manner by storing irreversibly masked data in semi-public storage and maintaining full encrypted data in a private repository.

Beyond blockchain, prior works have also explored specialized encryption schemes for genomics. Yamada et al. developed a system for genomic secret search using fully homomorphic encryption, enabling queries on encrypted DNA data without revealing the raw genome [30]. These earlier studies underscore the trend of marrying cryptography with genomic data sharing. However, many focused on clinical records or small genomic segments and did not fully tackle the unique scale and sensitivity of whole-genome data. While they demonstrate key concepts, from using blockchain for consent managementto homomorphic encryption for privacy, a gap remains in delivering an integrated solution that addresses genomics-specific challenges (massive file sizes, personal ownership of raw data, and long-term security needs).

The combination of blockchain and IPFS for decentralized data storage and management has been explored in other domains, including peer-to-peer file sharing, access control, and identity management. Early research into blockchain-IPFS integration focused on enhancing the efficiency of decentralized storage. Chen et al. proposed a peer-to-peer (P2P) file system architecture that enhances data availability and retrieval speed using IPFS [31]. Zheng et al. further optimized storage models for blockchain-integrated IPFS systems, demonstrating improvements in data indexing and integrity verification. Steichen et al. introduced a blockchain-based access control framework for IPFS, enabling decentralized policy enforcement [32]. Recent industry implementations, such as Microsoft ION, leverage blockchain and IPFS for self-sovereign identity (SSI), enabling decentralized and user-controlled authentication [33]. While these studies provide fundamental designs for blockchain-IPFS integration, they do not address the privacy and security concerns associated with genomic data storage, nor do they incorporate specialized cryptographic techniques for data confidentiality [34]. In contrast, our framework introduces a dynamic, smart contract-based consent mechanism that allows genomic data owners to grant, revoke, and audit access permissions in real time.

**Table 1:** Blockchain comparison to the previous system and our work

| System | Storage mode | Consent model | Revocation | Audit granularity | Crypto used | Implementation status |
| --- | --- | --- | --- | --- | --- | --- |
| ReGene | Off-chain data; on-chain pointers / integrity anchors | Generally static approvals (dynamic not emphasized) | Not explicitly detailed | Metadata-level / integrity logs | Symmetric + public-key (as reported) | Concept/design (prototype referenced) |
| LifeCODE.ai | Off-chain content with on-chain indexing | User-controlled sharing; dynamic not central | Policy-level mention (no benchmarks) | Traceability logs (coarse–moderate) | Symmetric + public-key (as reported) | Platform description / prototype |
| Park et al. | Off-chain data; on-chain control; local differential privacy (LDP) focus | Policy-driven; dynamic not primary focus | Not central / not reported | On-chain policy/logs (moderate) | LDP + conventional crypto | Concept/prototype |
| This work | Off-chain IPFS; on-chain ACL pointers | Dynamic, revocable consent via smart-contract ACLs | Yes (grant/revoke on-chain) | Event-level on-chain logs | AES (data), ECC (key encapsulation) | Working prototype (Ethereum Sepolia, MetaMask, Node.js, GUI) |

### Paper Roadmap

Our proposed framework introduces several novel features that significantly advance the state of the art in blockchain-IPFS solutions for genomic data sharing. In contrast to prior research, which often lacks specialized encryption or uses static access policies, our approach integrates new techniques to enhance security, flexibility, scalability, and compliance. The key innovative aspects are outlined below.

Hybrid AES-ECC Encryption: We design a tailored encryption scheme optimized for large genomic datasets. Unlike existing blockchain-IPFS approaches, which typically rely on a single encryption method, our framework leverages a hybrid of AES-256 and ECC to maximize security and efficiency. The genomic data itself is encrypted with AES-256, a symmetric cipher well-suited for bulk data encryption, ensuring that even whole-genome files are secured with minimal performance overhead. We then use ECC to manage encryption keys and access rights. ECC’s lightweight key pairs allow us to encrypt and distribute AES keys to authorized users with low computational costs and smaller key sizes. This dual-layer strategy ensures robust protection (via strong symmetric encryption) while enabling fine-grained, lightweight key management (via asymmetric ECC keys) – an approach not seen in earlier blockchain-IPFS frameworks focused on genomic data. By specifically tuning our encryption workflow for genomic sequences, the framework achieves higher confidentiality and performance than prior methods, which often did not account for the unique size and sensitivity of genomic information.

Smart Contract-Based Dynamic Consent Management: Our framework introduces a flexible and revocable access control mechanism using smart contracts, going beyond the static access control models of prior systems. In previous blockchain-based sharing models, once data access was granted to a party, it was typically fixed and difficult to revoke or modify. In contrast, we implement dynamic consent via Ethereum smart contracts (or an equivalent blockchain platform), which allows data owners to grant, monitor, and revoke access in real-time. Through a dedicated consent smart contract, a genomic data provider can specify who is allowed to decrypt or query their data, under what conditions, and can withdraw that permission whenever necessary. Any change in consent (e.g., a patient withdrawing permission for a researcher to use their genomic data) is automatically recorded on the blockchain and immediately enforced in the access control logic. This dynamic consent management is tailored to genomic privacy needs and provides a level of control and transparency unavailable in earlier static models. The use of smart contracts here ensures an immutable, auditable log of consent decisions and guarantees that only those with up-to-date permission (as defined by the latest state of the contract) can access the data. This is a significant improvement over prior approaches, enabling flexible, user-centric privacy management for genomic data sharing. On blockchains that support them, such as Ethereum, smart contracts (selfexecuting code stored onchain) can automate and secure datasharing agreements; other ledgers without smartcontract functionality may still serve as immutable logs, but our framework specifically targets a smartcontractcapable platform.

## II. Methods

The system architecture (Figure 1) consists of four main components: Ethereum blockchain, IPFS, and Hybrid Encryption Scheme. The Ethereum blockchain is the system’s backbone, providing a decentralized, immutable, and transparent platform for storing metadata, access control policies, and encrypted IPFS hashes. AES-256 in Galois/Counter Mode (GCM) is utilized for symmetric encryption of the genomic data files. ECC, specifically Curve25519, is employed for secure key exchange and digital signatures, ensuring that only authorized parties can access the encrypted data. Smart contracts play a crucial role in the framework by automating the enforcement of access control policies and maintaining a tamper-proof record of all access events. The smart contracts define the rules and permissions for granting and revoking access to genomic data, ensuring that only authorized users can access the files. Moreover, the smart contracts enable auditing capabilities, allowing for transparent data access and usage tracking. To assess the performance and scalability of the proposed framework, we conducted experiments using a high-performance server with the following specifications: (1) CPU: AMD Threadripper 3990X (64 cores, 128 threads), (2) RAM: 128 GB DDR4-3200, (3) Storage: 2 TB NVMe SSD, (4) OS: Ubuntu 20.04 LTS. The software environment consisted of: (1) Ethereum client:Geth v1.10.23, (2) IPFS: go-ipfs v0.12.2, (3) Programming languages: Node.js v14.17.0, Solidity 0.8.17.

**Figure 1.**
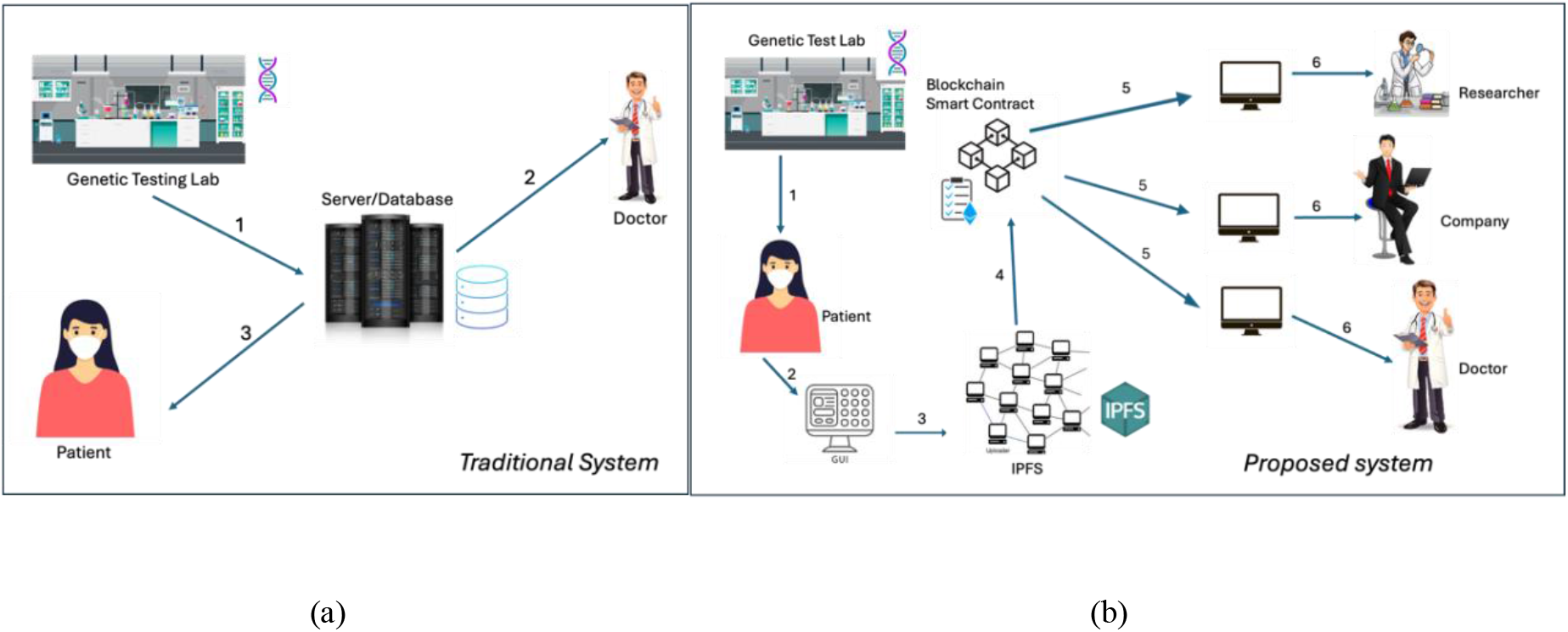
System Architecture for the traditional (a) and the proposed system (b)

### System Implementation on the Ethereum Sepolia Testnet

To validate the framework’s blockchain integration and data-sharing workflow, we deployed our system on the Ethereum Sepolia test network. The implementation involved a combination of tools and components, each serving a distinct role in the experimental setup:

MetaMask Wallet: We used the MetaMask browser wallet to manage Ethereum accounts and sign transactions on the Sepolia testnet. MetaMask facilitated user interactions with the blockchain by holding private keys for the data owner and authorized users and prompting transaction confirmations (e.g., when storing file metadata or updating access permissions on-chain). This ensured that all blockchain operations (such as deploying the smart contract, registering data, and granting access) were executed under real cryptographic signatures in a user-driven manner, closely emulating a practical deployment.Node.js Backend Script: The core backend logic was implemented in Node.js (v14.17.0). This script acted as the bridge between the system’s components, interfaced with the Ethereum blockchain (via a web3/ethers library connection to a Sepolia node) and with IPFS for file storage. The Node.js script handled the generation of cryptographic keys and the orchestration of data flow. For example, it generated the ECC key pair used for encrypting the AES key, and it formulated and submitted Ethereum transactions to the smart contract. All smart contract interactions (such as storing the IPFS content hash and encrypted AES key or updating the access control list) were executed through this Node.js layer. Using Node.js to interact with Sepolia, we ensured that the blockchain-related operations (smart contract calls, event listening, etc.) were carried out in a realistic network environment rather than a mock simulation.

3) Python GUI Application: A Python-based GUI was developed as the user interface for encryption and data handling tasks. This application allowed the data owner (e.g., “Sarah” in our use-case scenario) to easily encrypt genomic files and initiate the upload/share process without dealing directly with low-level commands. The GUI (running on Python 3.x) invoked the AES-256 encryption on the selected genomic data file and produced the ciphertext and a random initialization vector (IV). It then communicated with the Node.js backend (e.g., via HTTP requests or a local API) to pass along the encrypted file and trigger the subsequent steps (IPFS upload and blockchain registration). In essence, the Python GUI handled all encryption-related tasks on the client side and provided an intuitive front-end, while delegating blockchain and storage operations to the Node.js backend. Notably, Python was not used to simulate any blockchain behavior – it was confined to encryption, user interaction, and coordinating calls to the backend. This separation avoids any ambiguity: the blockchain logic ran on actual Ethereum infrastructure (through Node.js and Sepolia), not in a Python simulation.

4) Ethereum Sepolia Testnet and Smart Contract: We chose the Sepolia test network for deploying our Ethereum smart contract, which manages access control and metadata storage. Using a testnet provides a live blockchain environment (with real proof-of-authority consensus and transaction validation) without the cost and risk of mainnet deployment. We deployed a Solidity (v0.8.17) smart contract that contains functions to store the IPFS hash and encrypted AES key, and to grant or revoke access to specific Ethereum addresses. For the experiments, the contract was deployed and the relevant functions were called via transactions originating from the MetaMask-managed account of the data owner. Each transaction (e.g., registering a file’s hash/key, or updating access rights) was mined on Sepolia, producing a transaction hash and event logs as proof of execution. We verified these on-chain actions using Sepolia’s block explorer to ensure the correctness of operations (Figure 2b illustrates a sample transaction confirmation on Sepolia). The use of the Sepolia network in our blockchain-related experiments means that the integrity and behavior of our framework were tested under realistic conditions, including network latency and actual consensus, as opposed to a purely local simulation. At the same time, by using a testnet and modest data sizes for on-chain records (only hashes and keys), we kept the complexity and cost manageable.

**Figure 2.**
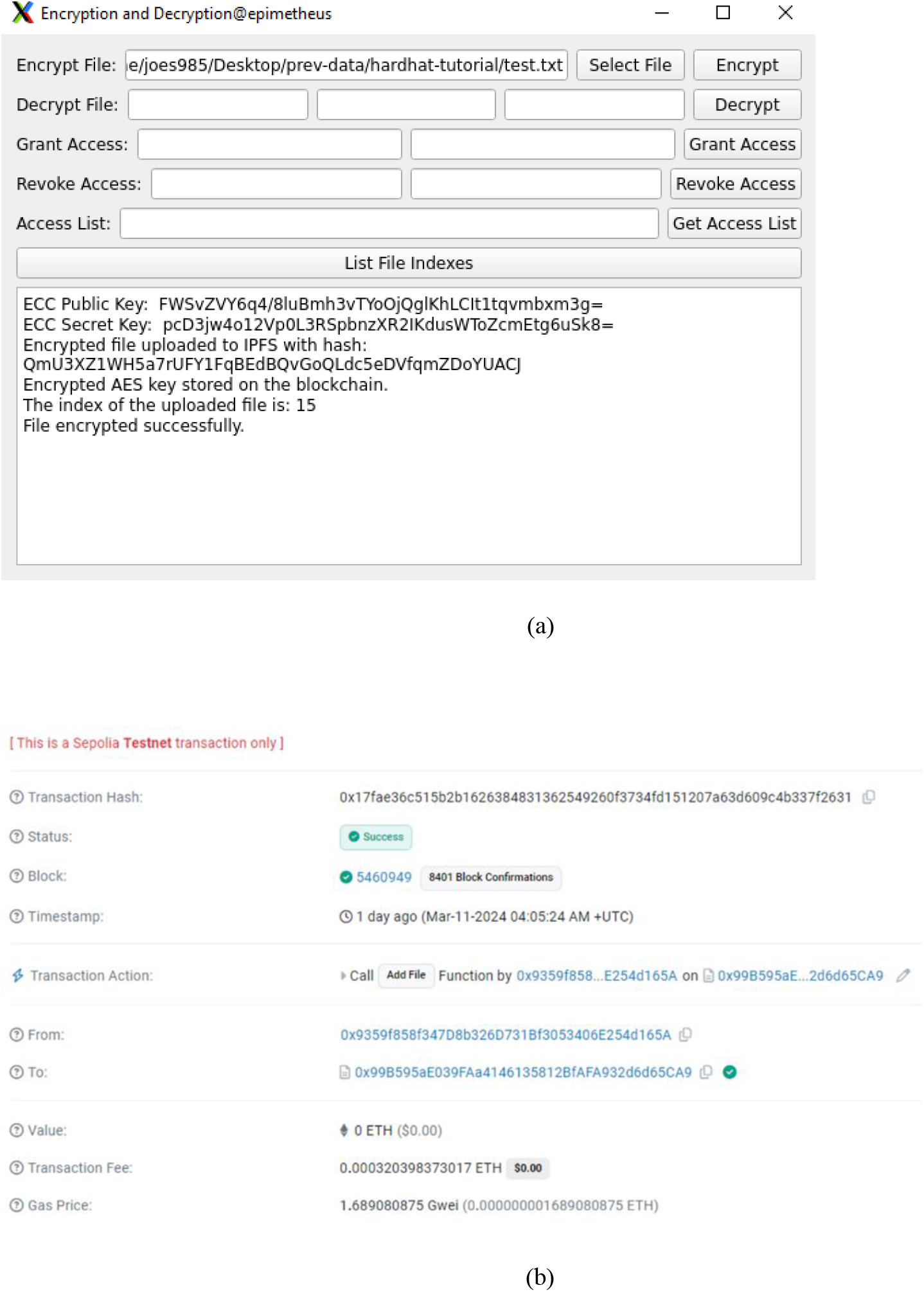
User interface. (a) The GUI interface used by the end-user (Sarah) for encryption. (b) Verification of the Transaction on the Sepolia Testnet.

5) IPFS: In our architecture, IPFS was the off-chain storage layer for the encrypted genomic data. The Node.js backend interacted with a local IPFS node (go-ipfs v0.12.2) to upload encrypted files and retrieve their content identifiers (CIDs). These CIDs (essentially content hashes) were then recorded on the blockchain via the smart contract, as described above. Integrating IPFS and Ethereum via the Node.js script provides an immutable linkage between the blockchain and the off-chain data.

Using the above setup, we performed end-to-end tests of the data-sharing workflow. First, the data owner used the Python GUI to encrypt a sample genomic dataset (the AES-256 symmetric key and IV were generated on the fly). Upon encryption, the GUI invoked the Node.js service to handle file upload and metadata registration. The Node.js script stored the encrypted file on IPFS (returning a CID) and generated an ECC key pair. The AES key from the encryption step was then encrypted with the recipient’s ECC public key (emulating a scenario where a specific researcher’s public key is used; in our test, a placeholder key pair represented the “authorized user”). Next, the script packaged the CID and the encrypted AES key into a transaction sent to the Ethereum smart contract on Sepolia (using the data owner’s MetaMask account for signing). This transaction added a new record on-chain linking the IPFS CID with its encryption key (now protected via ECC) and attributing it to the data owner’s address. We also tested the access control functions: the data owner sent a “grant access” transaction through MetaMask/Node.js to give the authorized user’s Ethereum address permission to download and decrypt that data. The smart contract updated its internal access control list and emitted an event, which the Node.js backend picked up to confirm the action. Finally, we simulated the data retrieval by having the authorized user’s client query the smart contract (via the Node.js script or a web3 call) to fetch the IPFS CID and encrypted AES key, then download the encrypted file from IPFS and decrypt it using the corresponding ECC private key and AES key. This step validated that a legitimate authorized party could retrieve and decrypt the data. Throughout these tests, MetaMask provided the transaction signing and approval interface, ensuring that all on-chain actions were user-authorized, while the Node.js backend manages blockchain and IPFS interactions. The Sepolia deployment confirmed that our framework’s blockchain components work in practice: we could immutably store data references on-chain, enforce dynamic access controls, and maintain the confidentiality of the genomic data, all in a decentralized environment. We did not measure blockchain performance metrics like throughput or latency, as our focus was on functional verification of the integration. In other words, the blockchain experiment was intended to demonstrate qualitative functionality (correctness of data storage and access control on a live network) rather than optimizing or quantifying the network’s performance.

To evaluate our framework without risking patient privacy, we used a combination of real reference genome data and synthetically generated DNA sequences. We first incorporated the publicly available human genome reference sequence (hg38), which provides an archetype of real genomic structure and length. In this study, we utilize hg38 (GRCh38), the latest human reference genome assembly, as a sample dataset for testing the functionality of our blockchain-based genomic data storage and access control system. The hg38 dataset was selected for: (1) High Data Complexity: The hg38 genome includes over 3.2 billion base pairs, making it an ideal test case to evaluate the system’s ability to handle large genomic files. (2) Public Availability: The reference genome is openly accessible, making it a suitable dataset for non-sensitive system validation without privacy concerns. In addition, we generated a diverse set of synthetic DNA sequences using a Python script. The goal was to simulate a range of genomic data scenarios – from small gene segments to whole-genome scales – in a controlled manner. Our synthetic data generator generated random nucleotide strings (comprising the bases A, T, G, and C) of varying lengths, designed to mimic the characteristics of real genomic data.We created sequences spanning a spectrum of sizes: some on the order of kilobases to represent individual genes or exons, and others on the order of hundreds of millions of bases to simulate entire chromosome-length sequences. This allowed us to assemble test datasets from a few megabytes to several gigabytes. In practice, we generated multiple datasets: for example, a small dataset (∼5 MB), comparable to a panel of genes, a medium dataset (∼500 MB), akin to an exome or a chunk of a genome, and a large dataset (several GB), approximating a whole-genome sequence. This range covers use cases from small-scale clinical genomics to large research cohort data. We chose to generate random sequences (as opposed to reusing someone’s real genome data) specifically to avoid any possibility of identifying an actual individual; the synthetic data has no relation to actual persons, yet it is similar in format and size to real genomic files. Using these methods, we ensured that our performance evaluation of the framework is relevant to real-world usage. The large synthetic genomes test the system’s scalability for throughput and storage (since they mimic the size of real genome files), which indicates how the framework would perform on actual genomic data in production.

In Figure 1, the traditional system, the genetic testing lab generates a patient’s genetic data and stores it on a central server or database (1). The doctor can access this data directly without the patient having any control or visibility over who is accessing it (2). The patient cannot regulate or manage who views her sensitive genetic information (3). Although she can access her data, she lacks transparency about who else may have accessed it and is not informed about any encryption methods (if any) used to protect her data. The security and privacy of the patient’s genetic information depend entirely on the centralized system, leaving her vulnerable to unauthorized access and unable to monitor or control data access. This highlights significant challenges regarding privacy and control in centralized databases, particularly when handling sensitive genetic information.

The proposed methodology has the following components: (1) Data Generation: The genetic lab generates the patient’s genetic data and sends it to her through a one-time use data source (like a jump drive). (2) Using a GUI interface, the genomic data encryption uses the AES algorithm with a 256-bit key via a mobile device or computer. A random initialization vector (IV) is generated for each encryption process to ensure uniqueness and security. (3) The encrypted genomic data is then uploaded to IPFS, a distributed storage system. Upon successful upload, IPFS returns a unique hash as an identifier for the encrypted genomic data. The metadata associated with the encrypted genomic data is prepared for storage on the Ethereum blockchain. (4) The metadata includes the IPFS hash, which refers to the encrypted data stored on IPFS. An ECC key pair, consisting of a public and private key, is generated by the Node.js script. The AES encryption key used to encrypt the genomic data is encrypted using the ECC public key. The encrypted AES key and other relevant metadata are then stored on the Ethereum blockchain via the smart contract. (5) The encrypted data is downloaded via an interactive GUI, which is downloaded by the respective parties (6).

A smart contract is deployed on the Ethereum blockchain to record and manage access control and data-sharing policies for genomic data. The smart contract contains the logic for granting and revoking access permissions to authorized third parties such as researchers, companies, and doctors. It also includes functions to verify and retrieve metadata stored on the blockchain. Third parties interested in accessing genomic data interact with the smart contract by submitting access requests, specifying: (1) the desired dataset (identified by an index) and (2) their Ethereum address for authentication. The smart contract verifies the requester’s permissions by checking the on-chain Access Control List (ACL). If authorized, the smart contract records the access event on the blockchain and allows the requester to retrieve the IPFS hash of the encrypted genomic file. State-changing access modifications (e.g., granting and revoking access) are logged on the Ethereum blockchain through event emission. The framework uses two key algorithms for handling genomic data. Algorithm 1: Encryption and secure storage of genomic data using AES-256 encryption before uploading to IPFS. Algorithm 2: Secure decryption and retrieval of genomic data via ECC-based key exchange and AES decryption.

### Encryption workflow

(1) Generate random AES-256 key and IV; (2) encrypt the genomic file with AES (client side); (3) encrypt the AES key using the recipient’s ECC public key; (4) upload ciphertext to IPFS and obtain CID; (5) write CID + encrypted AES key to the smart contract (access list).

### Decryption workflow

(1) Authorized user reads CID + encrypted AES key from the contract; (2) decrypt AES key with their ECC private key; (3) fetch ciphertext from IPFS; (4) decrypt file with AES. Note: ECC is used only for key encapsulation and signatures, not for bulk file encryption.

#### Algorithm 1

Genomic Data Encryption

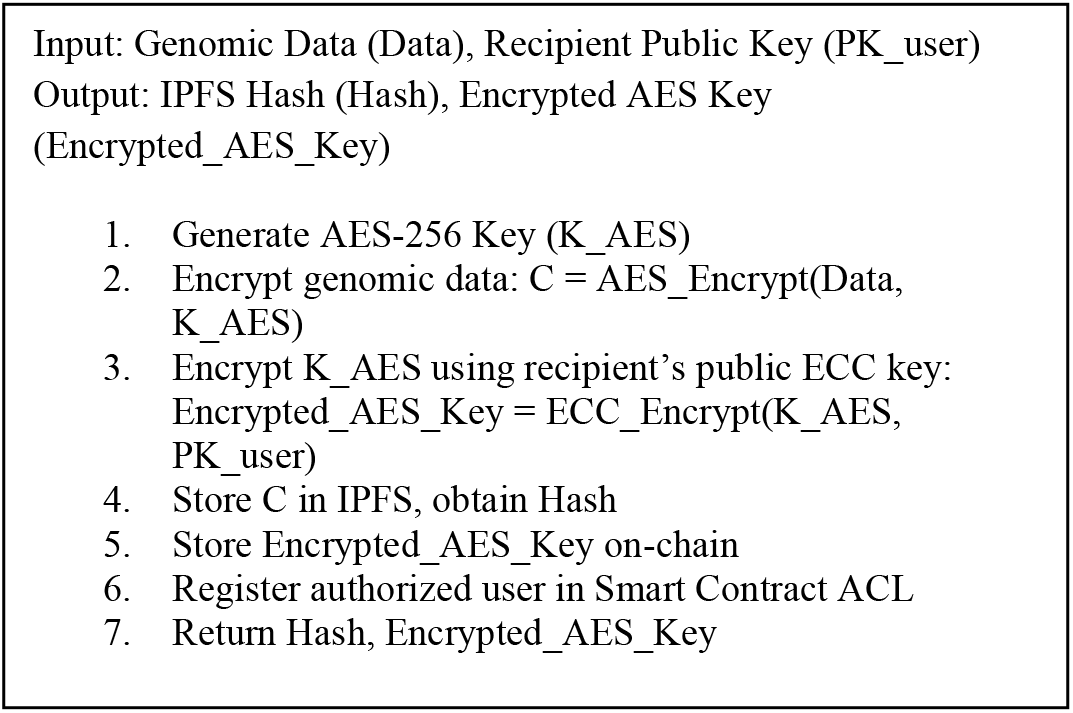

#### Algorithm 2

Genomic Data Decryption and Retrieval

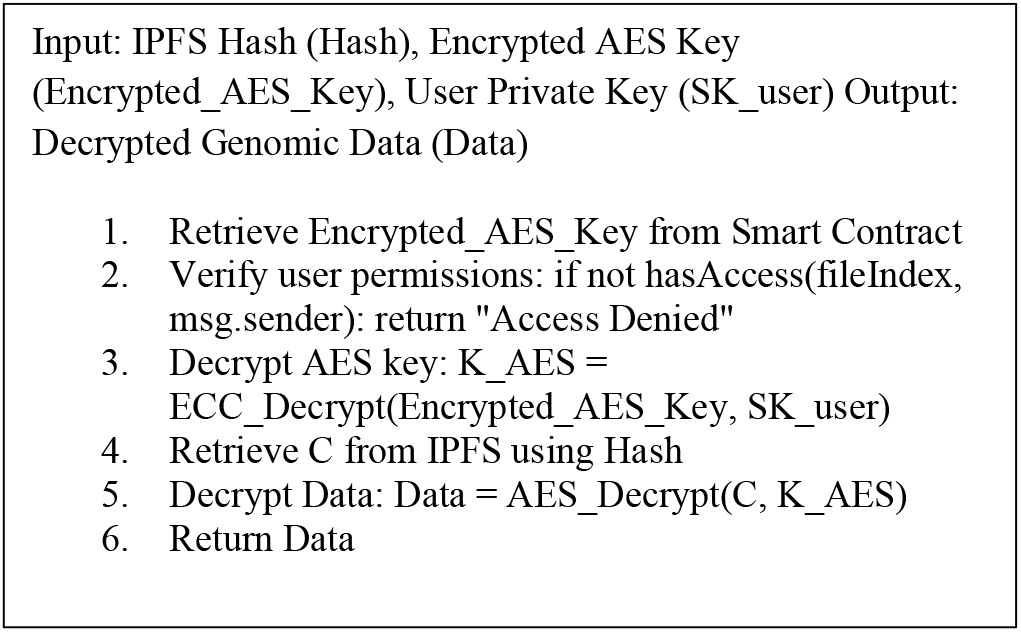

### Access Control Mechanism

#### Algorithm 3

Access Control

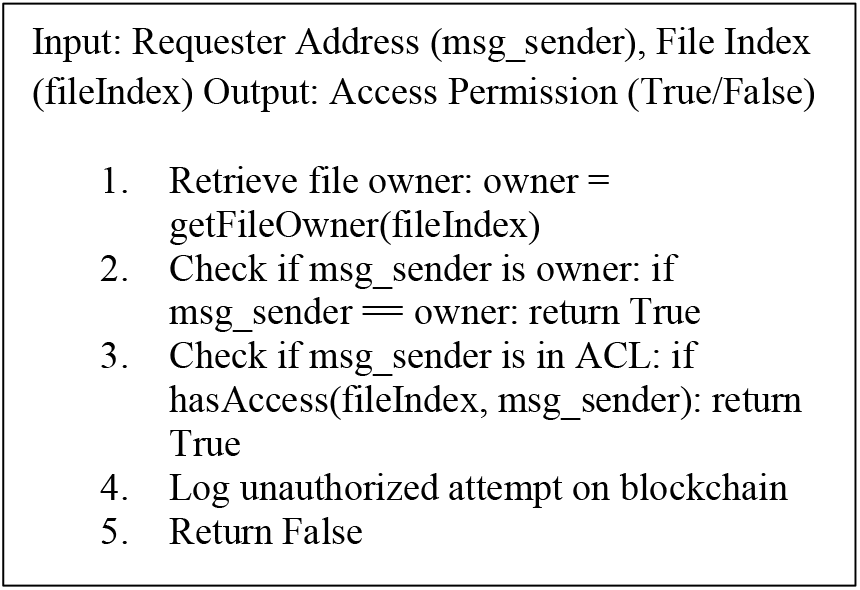

Unlike traditional access control systems that rely on centralized authentication mechanisms, our approach ensures that permissions are verifiable, tamper-resistant, and revocable in real-time. To enforce access control securely, we introduce a three-step verification mechanism that prevents unauthorized access to genomic data:

- Step 1: User Registration and Identity Verification
  ∘ Each genomic data owner and authorized user is identified through a unique blockchain address (msg.sender).
  ∘ When a new file is uploaded, the smart contract assigns ownership of the file to msg.sender, recording it in an ACL.
  ∘ Third-party users must request permission from the file owner before accessing the encrypted AES key.
- Step 2: Permission Verification and AES Key Retrieval
  ∘ When a user requests access, the smart contract executes the function:
  ∘ hasAccess(fileIndex, msg.sender)
  ∘ If the requesting user’s address is in the ACL, the smart contract releases the encrypted AES key.
  Otherwise, access is denied and logged on-chain to track unauthorized attempts.
- Step 3: Revocation and Dynamic Access Management
  ∘ The file owner can grant or revoke access at any time using:
  ∘ grantAccess(address_to_grant, fileIndex)
  ∘ revokeAccess(address_to_revoke, fileIndex
  ∘ Revocation removes the user from the ACL, immediately preventing further decryption.

### Key Management

A crucial aspect of our framework is secure key management. Each genomic data owner generates an AES-256 symmetric key for encryption. To protect this key, an ECC key pair is used, where the AES key is encrypted with the recipient’s ECC public key. This ensures that only authorized users can decrypt and access the genomic data. Key storage and distribution are handled securely within the blockchain-based system. The encrypted AES key and the corresponding IPFS hash are stored immutably on-chain. Only users registered in the smart contract’s ACL can retrieve and decrypt the AES key using their private ECC key.

Key revocation is a critical security feature that allows data owners to revoke access from a previously authorized user. This ensures that users without permission cannot decrypt or retrieve genomic data. The revocation mechanism operates through the following steps:

- Smart Contract-ACL Updates
  ∘ Each authorized user is registered via the blockchain-based ACL.
  ∘ When a data owner revokes access, the user’s Ethereum address is removed from the ACL, immediately preventing them from retrieving the encrypted AES key.
- Encrypted AES Key Invalidation
  ∘ Since the AES key is encrypted using the recipient’s ECC public key, revocation prevents the user from decrypting the AES key even if they previously accessed the encrypted file on IPFS.
- Immediate and Tamper-Proof Enforcement
  ∘ Once revoked, any future decryption attempts will fail, as the blockchain ensures that only authorized users can retrieve the AES key.
  ∘ The immutable nature of blockchain ensures that revoked access cannot be reversed without explicit permission from the data owner.

Architecture: While our current prototype stores the ciphertext of a wholegenome file as a single object on IPFS, this design indeed creates a “oneblob” focal point for an attacker. In this manuscript, we describe the end-to-end enforcement path (registration → ACL check → key retrieval → decryption) and demonstrate functional correctness on Sepolia. Quantitative measurements of access latency, concurrency, and revocation timing will be a future work. A more defenceindepth approach is to couple IPFS with a modular DA layer, e.g., EigenDA on EigenLayer or Celestia. DA networks first erasurecode each data blob (typically with 2D ReedSolomon) and then disperse the resulting shards across hundreds of restaked validator nodes; no single node retains the full file. To reconstruct the data, an adversary would have to compromise a statistically significant subset of nodes before they could even assemble the ciphertext, dramatically increasing the attack surface they must control. EigenDA, for example, uses a Disperser that posts a KZG commitment on-chain and economically slashes nodes that prove unavailable, while Celestia allows light clients to probabilistically verify that all shards remain online via data availability sampling. Integrating such a DA layer alongside IPFS would therefore add an orthogonal security layer, as well as geographic and economic dispersion, on top of cryptographic protection. We acknowledge this limitation of our current architecture and will explore an IPFS + DA hybrid in future work, particularly for ultra-sensitive datasets such as whole-genome sequences.

Validated the security claims: ProVerif is an automatic, symbolicmodel verifier that reasons soundly about an unbounded number of protocol sessions and message sizes. Version 2.04 (released 1 Dec 2021) introduced faster Hornclause saturation, native support for induction on natural numbers, and more aggressive trace pruning—features that shorten runtimes for complex, data-heavy workflows such as genome tokenization

While we recognize these limitations, they do not undermine the core concept of our framework. These issues require further work, optimizations, or complementary solutions. In the future, we plan to explore strategies such as integrating Layer-2 scaling solutions for blockchain, using decentralized storage incentives for IPFS, optimizing smart contract operations to reduce gas, and improving the UX of key management to address these challenges. Future work also includes the integration of standardized Post-Quantum Cryptography (PQC) schemes, such as CRYSTALS-Kyber, replacing ECC-based encryption to eliminate quantum vulnerabilities. Given current standardization timelines (e.g., NIST recommending a transition by 2030–2035), our architecture is designed to facilitate this critical upgrade with minimal effort and disruption.

### A Hypothetical Case Study

Sarah, a 35-year-old woman, recently underwent genetic testing at a leading healthcare facility to screen for the risk of breast cancer. The genetic testing process involved sequencing Sarah’s entire genome, which generated a vast amount of sensitive genetic information. After careful analysis of the sequenced data, the results revealed that Sarah carries a specific variant in the BRCA1 gene, which is known to be associated with a significantly increased risk of developing breast and ovarian cancer compared to the general population. Understanding the potential impact of her genetic information, Sarah recognized the importance of sharing her genomic data with breast cancer researchers. By contributing her data to research efforts, Sarah hoped to support the development of personalized medicine approaches and the identification of targeted therapies for individuals with similar genetic profiles. However, Sarah was deeply concerned about the privacy and security of her sensitive genetic information, given the potential risks of data breaches and unauthorized access in the digital age. To address Sarah’s concerns and provide a secure data-sharing solution, the genetic testing facility leveraged cutting-edge technologies to develop a robust and transparent data-sharing platform. The platform was powered by a Node.js script that seamlessly integrated three key components: IPFS, Ethereum blockchain, and a hybrid encryption scheme combining AES and ECC, as described in the following:

Sarah’s Process (Encrypting and Uploading Data)

- Generate AES-256 key and IV: Sarah’s application generates a secure 256-bit symmetric key (for AES-256 encryption) and a random Initialization Vector (IV) for encryption. This key (and IV) will be used to encrypt genomic data, providing confidentiality.
- Encrypt genomic data with AES-256: Using the generated AES-256 key, Sarah’s genomic data file is encrypted (e.g., with AES-256 in CBC or GCM mode using the IV). The result is ciphertext that appears as random data and cannot be understood without the same key.
- Upload encrypted data to IPFS and get CID: The encrypted genomic file (ciphertext) is uploaded to the IPFS, a distributed storage network. IPFS returns a CID, a cryptographic hash uniquely pointing to Sarah’s file on the network. This CID serves as a location-independent reference to the data.
- Generate an ephemeral ECC key pair: The system generates a fresh ECC key pair for the encryption session. This may involve creating a one-time (ephemeral) ECC private key and a corresponding public key. The ECC keys will be used to securely share the AES key with the intended recipient. In practice, Sarah already possesses an ECC key pair (public/private) for the platform, and the recipient (Dr. Greene) has his ECC public key available. An ephemeral key is used internally by the encryption algorithm to ensure forward secrecy during this process.

5. Encrypt the AES key with ECC (recipient’s public key): Sarah’s 256-bit AES key is then encrypted using ECC public-key encryption so that Dr. Greene can later decrypt it. Specifically, the system uses Dr. Greene’s ECC public key to encrypt the AES key (often via an ECIES scheme, which under the hood uses an ephemeral ECC key and Dr. Greene’s public key to derive a shared secret). This produces an encrypted AES key blob that only Dr. Greene (with his corresponding ECC private key) can decrypt. Encrypting the symmetric key ensures that even if someone obtains the key record, they cannot read the actual AES key without the proper ECC private key.

6. Store IPFS CID and encrypted AES key on Ethereum: Sarah interacts with an Ethereum smart contract to register the file’s metadata on the blockchain. She sends a transaction to the smart contract that stores the IPFS CID (content hash of the encrypted file) and the encrypted AES key (the small cipher blob from step 5) on-chainThe actual genomic data itself is not stored on Ethereum – only the reference (CID) and the encrypted key are kept on-chain, which provides a tamper-proof record of the data’s location and the key needed to decrypt it (in protected form). This on-chain record is immutable and auditable while revealing no usable genomic data publicly. For data availability, pinning is performed by the deploying provider, with optional co-pinning by the data owner via the client. A managed pinning service may be used for redundancy; only content identifiers (CIDs) are recorded on-chain.

Grant access to Dr. Greene via Ethereum transaction: Using another Ethereum transaction, Sarah calls a smart contract function (e.g., grantAccess) to give Dr. Greene permission to access that specific genomic dataset. She provides Dr. Greene’s Ethereum address (which is linked to his identity on the platform) and the identifier for the data (or its index/CID) in the transactionThe smart contract adds Dr. Greene’s address to an ACL for that file, recording that he is an authorized userThis permission is recorded on-chain, and the transaction emits an event logging that Dr. Greene was granted access. From this point, the blockchain stores: (a) the file’s IPFS CID, (b) the encrypted AES key, and (c) the list of addresses (including Dr. Greene’s) allowed to retrieve the key and access the data.

Dr. Greene’s Process (Retrieving and Decrypting Data)

- Verify on-chain access permission: Before attempting to fetch any data, Dr. Greene (the researcher) must be authorized. His client (or DApp) checks the Ethereum smart contract’s ACL to confirm that his Ethereum address has permission to access Sarah’s genomic dataset. This could be done by calling a read function (e.g., hasAccess) on the contract to see if Dr. Greene’s address is listed for the file. If he is not authorized, the system will deny access immediately (and this attempt can be logged on-chain). Only if the blockchain indicates that Dr. Greene’s address has been granted access will the retrieval process proceed.
- Retrieve IPFS CID and encrypted AES key from the blockchain: Once permission is confirmed, Dr. Greene (or the application on his behalf) retrieves the file’s metadata from the Ethereum smart contract. This includes fetching the IPFS Content ID (the hash pointer to the encrypted file) and the encrypted AES key associated with the file, which Sarah stored on-chain earlier. These pieces of information are typically stored in the contract’s state (often indexed by a file ID or hash). By reading the blockchain data, Dr. Greene obtains the CID (needed to download the file from IPFS) and the small encrypted symmetric key blob. (In many implementations, the act of retrieving this metadata can also trigger a blockchain event recording that Dr. Greene has requested access to at this time
- Decrypt the AES key using ECC private key: Dr. Greene now uses his ECC private key to decrypt the symmetric AES key. Using his private key and the encrypted key blob from the blockchain, he performs an ECC decryption, which recovers the original 256-bit AES key that Sarah had used to encrypt the dataThis step relies on Dr. Greene’s private key matching the public key that Sarah used in step 5 of her process. Thanks to the asymmetric encryption, only Dr. Greene can obtain the plaintext AES key; no one else who might see the encrypted key (on-chain) can derive it without Dr. Greene’s private key.
- Fetch the encrypted file from IPFS: Armed with the IPFS CID, Dr. Greene’s application connects to the IPFS network and downloads the encrypted genomic file. Using the CID ensures he gets exactly the data that Sarah uploaded (IPFS will retrieve the correct content-addressed file from the network). The file he downloads is still in encrypted form (the AES-encrypted binary), so even if someone intercepted this IPFS transfer, it would be unreadable without the AES key.
- Decrypt the genomic data with AES-256: Dr. Greene uses the recovered AES key (and the original IV, which may be provided alongside the key or derived) to decrypt the genomic data file using AES-256. The AES decryption transforms the ciphertext back into the original genomic data in plain form, allowing Dr. Greene to view or analyze the genome information. At this point, Dr. Greene has full access to the sensitive data, which was protected until the decryption. The successful access can be recorded (if the system logs the event on-chain or off-chain) for audit purposes.

### Access Control and Audit Logs

On-chain permission management (ACL): The Ethereum smart contract functions as the access control layer. It maintains an Access Control List mapping each encrypted genomic file (often referenced by a file index or CID) to the list of authorized Ethereum addresses. Data owners like Sarah can call contract functions such as grantAccess(address, fileId) or revokeAccess(address, fileId) to update these permissions at any time. Granting access adds a user to the allowed list for that dataset, while revoking access removes them, immediately preventing any further decryption by that address. These functions are executed via Ethereum transactions, ensuring that only the file owner (or an authorized delegate) can modify who has access. The ACL check (hasAccess) is enforced every time a user tries to retrieve a key or data, so only addresses in the allowed list can proceed.

Grant/revoke transparency: Every change in permissions is transparently logged on the blockchain. When Sarah grants access to Dr. Greene or later revokes it, the smart contract emits events that record the action.

Logging data access attempts: The system also logs data retrieval events for auditing. Whenever Dr. Greene (or any researcher) attempts to access the data, the smart contract can record an event. If the access is authorized, the contract can emit an event noting that Dr. Greene accessed file X at time Y. If an unauthorized address tries to access the data (or if Dr. Greene tries to access a file he isn’t allowed to), the contract’s permission check fails and it can emit a security event or log the attempt as “access denied” with the user’s address and timestampThese events create a permanent audit trail of both successful and attempted data accesses.

Tamper-proof audit trail: Because all grants, revocations, and access events are recorded on Ethereum’s distributed ledger, neither Sarah nor Dr. Greene (nor any malicious actor) can alter the history of who did what. The audit log is tamper-proof and can be inspected by authorized parties at any time to verify compliance and usage patterns. Figure 2a shows the GUI interface and Figure 2b shows the backend processing on the Sepolia Testnet.

Post disclosure ownership and downstream duties: Legal title to the genomic record remains with the patient under HIPAA (45 CFR §164.502(g)). The physician could receive only a timebounded AES key released by the smart contract; once the key expires (default 72 h) or the patient issues revokeAccess, the ciphertext is no longer decryptable. Any derivative artefacts (VCF files, clinical notes, and risk scores) generated during the authorized window must embed the originating transaction hash in their metadata. Institutional policy, enforced by audit, then requires deletion or reauthorization of those artefacts once the key is invalid. Thus, even after the data leaves the onchain enclave, the patient retains enforceable control over further processing, storage, and reuse of their genome.

### Regulatory Compliance Considerations

Genomic data is highly sensitive personal information, and any management system must adhere to privacy regulations and ethical standards. Our proposed framework was designed with regulations like the European GDPR in mind, as well as the need for robust patient consent management. We discuss here how the system aligns with these regulations and what measures are taken to ensure compliance.

#### General Data Protection Regulation (GDPR)

The GDPR imposes strict requirements on handling the personal data of EU citizens, including genomic data. One key aspect is the right to be forgotten, which allows individuals to request the deletion of their data. On a surface level, blockchain’s immutability seems at odds with this requirement, since data on-chain cannot be altered or removed. However, our framework avoids storing any raw personal genomic data on-chain. The blockchain only holds encrypted keys and content hashes, which do not identify an individual or reveal genomic information in isolation. The actual genomic files reside in IPFS, which can be deleted or made inaccessible if a patient wishes to withdraw consent. To comply with a deletion request, a patient could ask for their IPFS-hosted file to be removed (or simply cease pinning it so that it is no longer distributed) and destroy their encryption keys, rendering the data effectively unreadable and thus “forgotten” in practical terms. The on-chain record (a hash pointer) is anonymous and contains no personal identifiers; it remains only as a historical ledger entry, much like a receipt, without violating privacy. This approach of detaching personal data from the ledger is in line with recommendations to reconcile blockchain with GDPR. Additionally, GDPR mandates transparency and limits the purpose of data use. Through our smart contract, every access to data is transparently logged, and patients could be informed or even required to approve each access, thereby upholding principles of informed consent and purpose limitation. Since those events are immutably recorded, our system can facilitate audits to show regulators which entities accessed data and for what stated purpose. Finally, we ensure compliance with GDPR’s data minimization principle. Only the minimal necessary information (essentially a pointer and an encrypted key) is handled by the blockchain, greatly reducing the risk profile if the ledger itself is analyzed by an adversary. By combining off-chain data storage with on-chain tokenization, we believe the framework can meet GDPR requirements and protect individual rights while benefiting from blockchain features. Our framework separates content and metadata to support data minimization: genomic files remain off-chain and encrypted, while on-chain records contain only pseudonymous references (CIDs) and policy state. We note that, under evolving EU guidance, even such identifiers or logs could be interpreted as personal data depending on context. To align with GDPR principles, we support: Right to be forgotten: key revocation and unpinning of off-chain files, combined with on-chain de-referencing, effectively prevents further access. Purpose limitation: audit logs record only access events related to genomic data sharing; no secondary-use attributes are written.

Record minimization: event schemas are limited to timestamps, pseudonymous addresses, and policy state. Accountability: immutable on-chain logs provide verifiable evidence of consent changes and access actions.

### Patient Consent and Ethical Data Use

Beyond formal regulations, ethical genomic data management requires respecting patient consent and preferences. Our blockchain-based design inherently supports a dynamic consent model. Patients (data owners) hold the keys to their data and can smart-contractually grant or revoke access to others in a granular way. For example, a patient could allow a university’s medical research team to access their genomic data for a specific study. This consent would be represented on the blockchain (either implicitly by sharing the key or explicitly by a consent record), and the patient could later revoke it, cutting off the research team’s future access. Such a mechanism aligns with modern approaches to genetic data consent, emphasizing participant empowerment and ongoing control. The blockchain would then record this consent transaction, and only then would the researcher obtain the decryption key. Such a workflow ensures compliance not only with legal standards like GDPR but also with ethical best practices, by making data sharing transparent and revocable. Another regulatory aspect is ensuring that our system does not inadvertently enable the misuse of data. Because genomic information can reveal familial relationships, ancestry, and health risks, our access control must prevent unauthorized sharing or public posting of someone’s genome. The legal responsibility in many jurisdictions is on data custodians to avoid breaches. Under our model, each data access is deliberate and logged, which creates accountability – a researcher cannot plausibly claim an accidental data leak without it being evident how they accessed the data. Furthermore, by using cryptographic tokens, we can integrate with emerging self-sovereign identity frameworks where patients verify the identity of requesters before granting access, satisfying requirements for informed consent documentation and verification of data recipients.

### Security Analysis

Even with strong encryption and blockchain integration, it is crucial to analyze the security of each component of the framework. We consider smart contracts controlling access, the IPFS off-chain storage, and the cryptographic schemes, discussing potential vulnerabilities and mitigation strategies.

#### Smart Contract Security

The Ethereum smart contract in our system mediates access to genomic data by enforcing permissions and logging access events. Because smart contracts are immutable programs that handle valuable permissions, they are a potential target for attacks. A known class of vulnerabilities in Ethereum contracts (reentrancy attacks, arithmetic overflows, or unauthorized access flaws) could be exploited if the contract is not carefully designed. We mitigated these risks by following established secure coding practices. The contract’s functions for granting or revoking access are structured to avoid complex state changes after external calls, thereby preventing reentrancy issues. We also employ Solidity’s overflow-safe math libraries (or use built-in overflow checks) to avert numeric errors. Additionally, the contract is kept as simple as possible: its primary roles are to store metadata (encrypted keys and IPFS CIDs) and to check signatures/permissions. Every access request is authenticated by requiring the requester to call the contract using their own Ethereum address, which is bound to their cryptographic identity; thus, the contract can verify permissions without handling plaintext credentials. We conducted thorough testing of the contract, including unit tests and testnet deployments, to validate that only authorized users can trigger the release of an encrypted AES key. By taking these precautions, the smart contract provides a robust and tamper-resistant access control layer. However, it is important to note that no smart contract is entirely immune to attacks — ongoing updates and security audits will remain necessary as potential new vulnerabilities are discovered in the Ethereum ecosystem.

#### Ipfs Data Security

Using IPFS for genomic data storage introduces its own security considerations. IPFS is a distributed file system where files are addressed by the hash of their content using CID. A major benefit of this content-addressing is data integrity: any modification to the file would produce a different hash, so IPFS inherently prevents undetected tampering — if someone altered even a single nucleotide in the encrypted file, the CID stored on the blockchain would no longer match, alerting that the content had changed or been corrupted. This gives our framework a built-in way to verify that the off-chain data retrieved is precisely what was originally uploaded. However, IPFS does not provide encryption or access control on its own; by design, IPFS content is accessible to anyone who knows the content hash. In our system, this is acceptable because all genomic files on IPFS are already encrypted – even if an unauthorized party obtained the CID and downloaded the file, they would only see ciphertext. This underscores the importance of strong encryption and key secrecy. We ensure that the AES encryption keys are never stored in IPFS, only on the blockchain in encrypted form and released to authenticated users. Developers using IPFS for sensitive data typically must add encryption at the application layer, since IPFS by itself stores data in plain form. Our approach follows this best practice by encrypting all genomic data before IPFS storage.

One potential vulnerability in IPFS usage is data availability. While IPFS is distributed, content is not guaranteed to be stored forever by the network unless a node pins it. In our deployment, either the data owner or a trusted service pins the encrypted genomic data to ensure it remains accessible. If only one node stores the data and goes offline, it may become temporarily unavailable (even though the blockchain reference remains). We address this by redundantly pinning data on multiple IPFS nodes (for example, run by the hospital, research institution, or third-party pinning services). This redundancy mitigates the risk of data loss or downtime. Another consideration is that an attacker could attempt a denial of service by requesting content from IPFS repeatedly or trying to overload the network. Such attacks are non-trivial to aim specifically at our content, since IPFS uses a swarm of nodes and content addressing to distribute load but monitoring and rate-limiting access requests (possibly via the smart contract or an off-chain proxy) can provide additional defense. Overall, by combining IPFS with encryption and proper pinning strategies, the system ensures data integrity and confidentiality. The blockchain’s immutable record of the CID means that even if a malicious actor tried to inject a fake file into IPFS, they could not alter the true CID on the ledger, and users would detect the mismatch immediately. Thus, the IPFS off-chain storage, when used in tandem with our on-chain controls, remains secure and reliable for our purposes.

The present security analysis establishes that access control is enforced by a smart contract, data remains encrypted end-to-end with an AES key protected under ECC, and payloads are stored off-chain via IPFS with only minimal, pseudonymous references recorded on-chain. While these properties substantially reduce the likelihood of a direct cryptographic break, sophisticated adversaries typically bypass protocols rather than break them. Accordingly, we extend the threat model to include social engineering and key theft, endpoint compromise, insider abuse and collusion in the storage and data-availability layer, and supply-chain manipulation of client software. The objective is not merely to enumerate attacks but to integrate layered, verifiable defenses that align with our existing revoke/expiry mechanisms and privacy posture.

First, social-engineering campaigns that capture credentials or wallet control present a direct path to lawful-looking misuse: an attacker who compromises the Ethereum identity authorized by the contract can legitimately trigger a key release without touching the cryptography. To address this, we require phishing-resistant authentication (e.g., FIDO2/WebAuthn) for all administrative consoles, mandate hardware wallets or HSM-backed custody for keys that can authorize decryptions, and use out-of-band confirmations for sensitive on-chain actions such as grant and revoke. Threshold approvals for releasing wrapped AES keys (for example, two-of-three among patient, institution, and a designated recovery guardian) further eliminate single-key failure. These controls complement the existing time-bounded key design and ensure that revocations take effect promptly even under active phishing conditions.

Second, endpoint compromise remains a critical risk because plaintext necessarily exists at the point of use. Our mitigation strategy seals the ECC-wrapped AES key to a trusted execution environment and performs unwrap and decrypt inside an attested enclave, emitting verifiable attestation evidence alongside the access event. Decrypted outputs are delivered only to ephemeral workspaces that automatically wipe on session end and enforce clipboard and disk-write guards, reducing the surface for memory scraping and post-exfiltration abuse. Because these steps integrate with the same on-chain access policy and revocation logic, they do not require changes to the contract interface while materially strengthening the operational path where data are most vulnerable.

Third, availability and confidentiality risks persist within the storage layer. IPFS and pinning services enhance integrity through content addressing; however, single-provider capture, insider threats, selective withholding, or collusion can compromise availability or leak side information through access patterns. We therefore distribute content across multiple independent pinning providers and institutional nodes, optionally erasure-code replicas to raise the collusion threshold, and run periodic proof-of-availability sampling with alerts for unexpected unpinning. These measures coexist with our GDPR-aligned “unpin plus key destruction” workflow: while unpinning is essential for honoring consent withdrawal and data minimization, independent monitoring ensures that an adversary cannot quietly mimic such actions to censor or coerce.

## III. Results

We conducted experiments to evaluate the performance of the Hybrid AES/ECC (HEC) protocol in terms of encryption time, decryption time, and data processing throughput. The experiments were conducted on a Linux machine using Python, as the hardware environment utilized on a server comprised of an AMD Treaddripper 64-core with 128 GB RAM.

Figure 3 compares encryption (En) and decryption (De) time, measured in microseconds (µs). Hybrid AES/ECC provides extra security with virtually no additional time in the encryption scheme and less time in the decryption scheme. The comparative analysis of encryption times serves to evaluate our proposed Hybrid AES/ECC (HEC) protocol against established encryption methods. We compared HEC with several widely used algorithms: AES, known for its efficiency and strong security; Blowfish, a symmetric block cipher with a variable key length; TripleDES (Triple Data Encryption Standard), which applies the DES cipher algorithm three times to each data block; and AES/RSA, a hybrid approach combining AES with the RSA public-key cryptosystem. Our hypothesis was that HEC would offer a superior balance of security and efficiency. The results for the smallest data size of 100 KB support this hypothesis: HEC (represented as AES/ECC in our data) achieves an encryption time of 170.71 µs, outperforming standard AES (286.1 µs), Blowfish (796.08 µs), TripleDES (2819.3 µs), and AES/RSA (165.22 µs). This performance advantage is particularly significant given that HEC combines the strengths of both AES and ECC, potentially offering enhanced security without compromising efficiency. These findings suggest that HEC could be valuable in scenarios requiring both high security and computational efficiency, such as in resource-constrained IoT devices or high-throughput data processing systems. The performance trends across various data sizes, as illustrated in Figures 3 and 4, further reinforce HEC’s potential as a robust and efficient encryption solution for modern cryptographic challenges.

**Figure 3.**
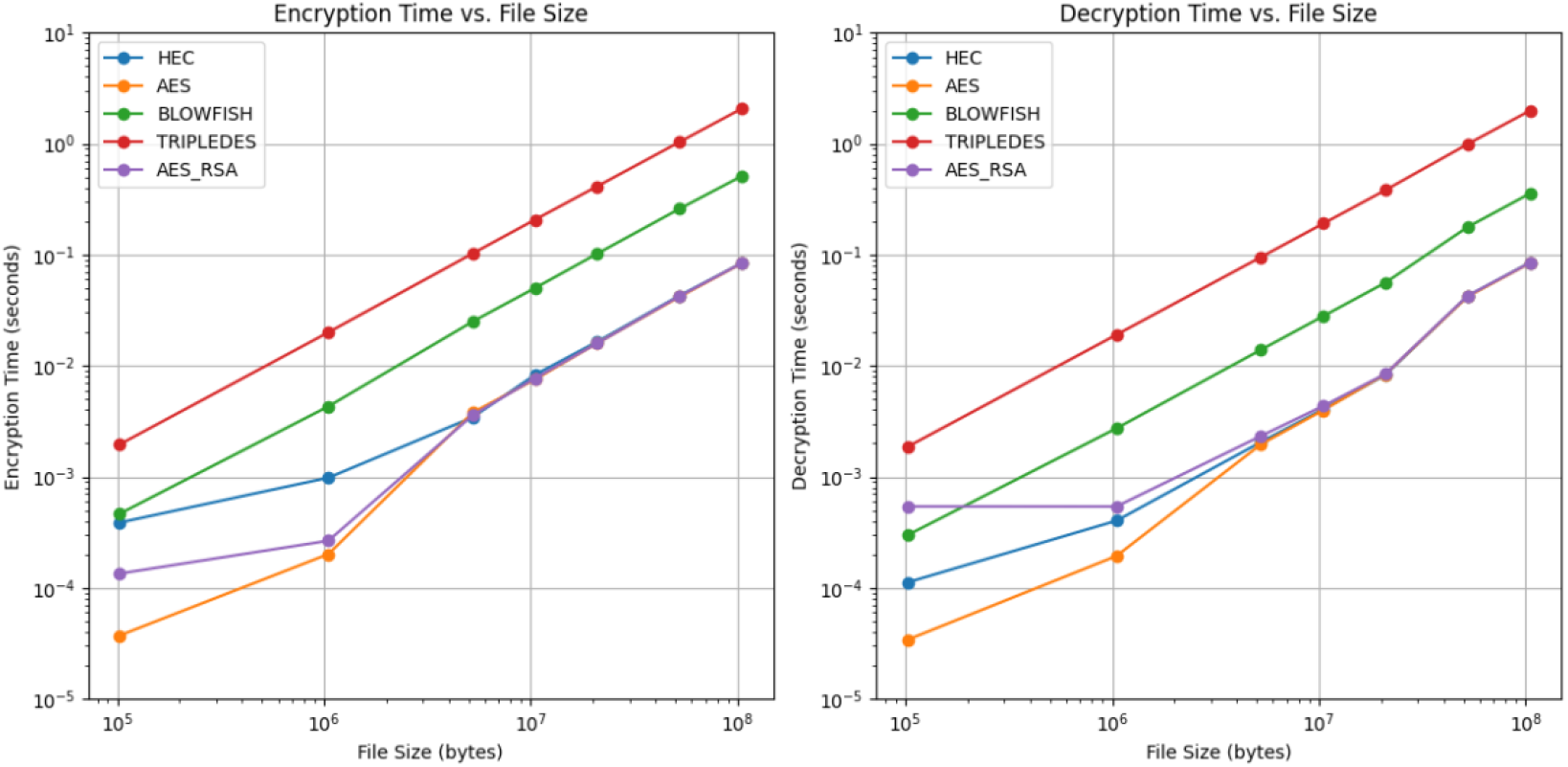
Encryption and decryption time (microseconds, µs) versus data size (KB, MB) for different encryption schemes: Hybrid AES/ECC (labeled as ECC/AES), AES, Blowfish, TripleDES, AES/RSA. X-axis: Data Size (KB, MB); Y-axis: Time (µs).”

**Figure 4.**
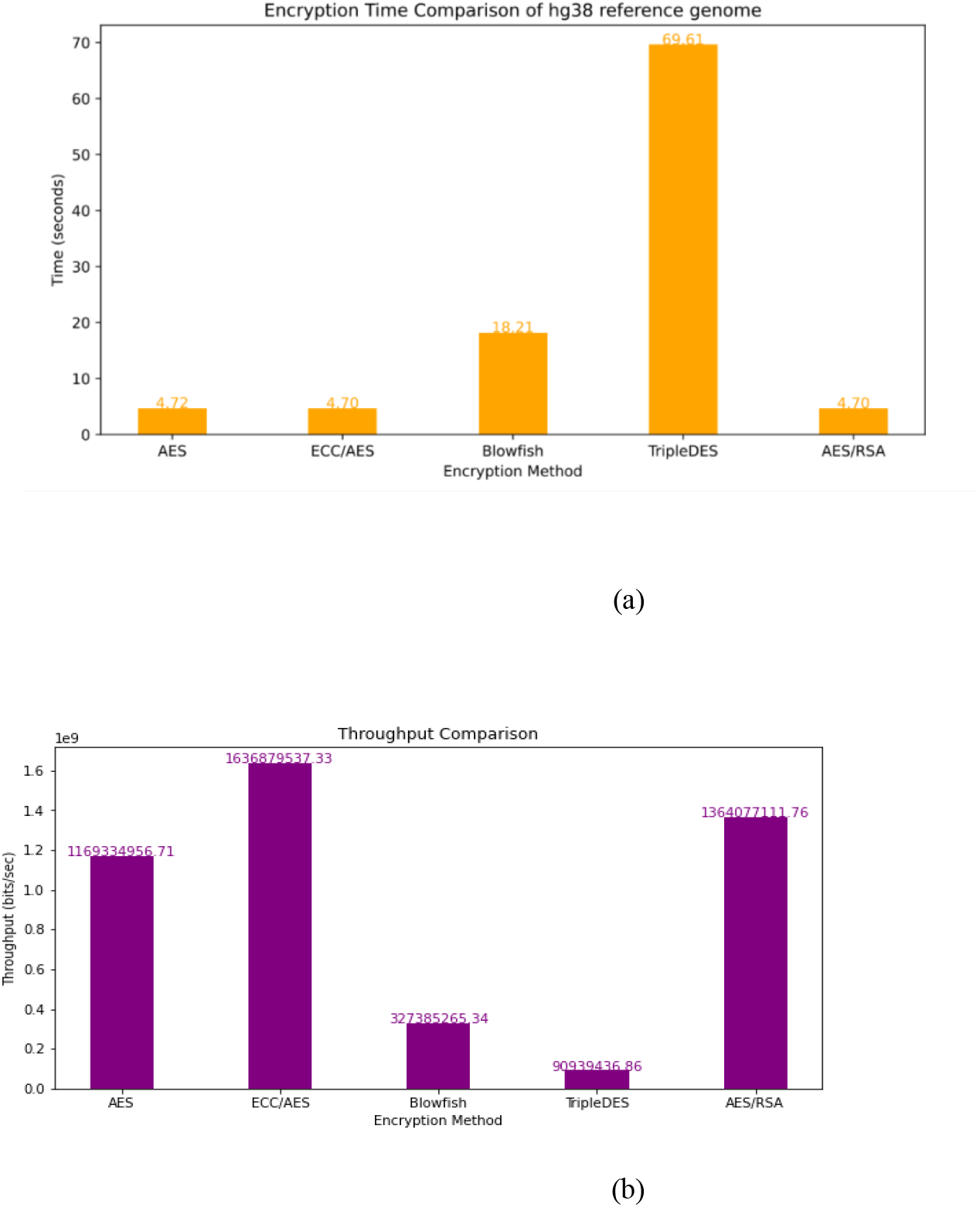
(a) Total encryption time (seconds) for the hg38 reference genome using different algorithms (AES, ECC/AES, Blowfish, TripleDES, AES/RSA). (b) Measured throughput (MB/sec). X-axis: Algorithm; Y-axis: Time (seconds) or Throughput (MB/sec).

It is worth noting that the performance difference between AES and AES/ECC is relatively small. For example, encrypting 100 MB of data takes 159.26 µs with AES, while AES/ECC requires 175.7 µs, resulting in a difference of 16.49 µs in favor of AES. Consistent with Figure 4a, AES is the fastest for symmetric bulk encryption. The hybrid AES/ECC design uses AES only for data-plane encryption and ECC for key distribution; therefore, we do not claim that the hybrid outperforms AES in data-plane speed. The performance of Blowfish is noteworthy, as it consistently outperforms TripleDES and AES/RSA for encryption and decryption across all data sizes. For instance, when encrypting 50 MB of data, Blowfish takes 84.8 µs, while TripleDES and AES/RSA take 1415.73 µs and 84.16 µs, respectively. Similarly, for decrypting 50 MB of data, Blowfish takes 385.76 µs, while TripleDES and AES/RSA take 1414.5 µs and 81.0 µs, respectively. TripleDES, despite its widespread adoption, exhibits significantly slower performance than the other algorithms. Its encryption and decryption times are consistently higher across all data sizes. For example, encrypting 10 MB of data takes 295.8 µs with TripleDES, while AES, AES/ECC, Blowfish, and AES/RSA take 31.71 µs, 29.0 µs, 109.4 µs, and 29.56 µs, respectively. Similarly, decrypting 10 MB of data takes 291.3 µs with TripleDES, while AES, AES/ECC, Blowfish, and AES/RSA take 25.0 µs, 25.1 µs, 107.0 µs, and 25.03 µs, respectively. AES/RSA, which combines the strengths of AES and RSA, exhibits the slowest performance among the compared algorithms. The overhead the RSA component introduces results in higher encryption and decryption times, especially for larger data sizes. For instance, encrypting 100 MB of data takes 159.4 µs with AES/RSA, while AES, AES/ECC, Blowfish, and TripleDES take 159.2 µs, 175.7 µs, 768.9 µs, and 2818.4 µs, respectively. Similarly, decrypting 100 MB of data takes 149.4 µs with AES/RSA, while AES, AES/ECC, Blowfish, and TripleDES take 149.4 µs, 138.9 µs, 2808.3 µs, and 4308.8 µs, respectively.

Figure 4a illustrates the encryption time comparison for the hg38 reference genome. TripleDES has the highest encryption time at 69.61 seconds, indicating it is the slowest among the compared algorithms. AES/RSA follows with an encryption time of 3.20 seconds. Blowfish and AES/ECC have similar encryption times of 18.21 seconds and 3.70 seconds, respectively. AES exhibits the lowest encryption time at 2.72 seconds, making it the fastest algorithm for encrypting the hg38 reference genome. Figure 4b presents the throughput comparison, which measures the amount of data processed per unit of time. AES demonstrates the highest throughput at 1634.7 MB/sec, confirming its efficiency in encrypting large datasets. AES/ECC follows closely with a throughput of 1198.9 MB/sec. Blowfish and AES/RSA have throughputs of 243.3 MB/sec and 1384.0 MB/sec, respectively. TripleDES has the lowest throughput at 63.5 MB/sec, aligning with its longer encryption time.

The avalanche effect is a desirable property in cryptography that refers to the sensitivity of the output (ciphertext) to small changes in the input (plaintext or key). In other words, when a single bit in the plaintext or key is changed, it should result in a significant and unpredictable change in the ciphertext. The avalanche effect is crucial for the security of the encryption algorithm because it makes it difficult for an attacker to deduce any meaningful information about the plaintext or key by analyzing the ciphertext. It is closely related to diffusion, which spreads the influence of each plaintext or key bit across many ciphertext bits, making the relationship between the plaintext, key, and ciphertext as complex as possible. The avalanche effect can be quantified using the avalanche effect score or the avalanche criterion. The avalanche effect score measures the average number of bits that change in the ciphertext when a single bit is flipped in the plaintext or key. The ideal avalanche effect score is 0.5, meaning that, on average, half of the ciphertext bits should change when a single bit is modified in the input. The equation for calculating the avalanche effect score is as follows in Equation 3:

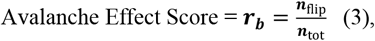

In practice, the avalanche effect score is calculated by performing the following steps:

- Encrypt the original plaintext with the original key to obtain the original ciphertext.
- Flip a single bit in the plaintext or key.
- Encrypt the modified plaintext with the modified key to obtain the new ciphertext.
- Compare the original and new ciphertext bit by bit and count the number of flipped bits.
- Divide the number of flipped bits by the total number of bits in the ciphertext to obtain the avalanche effect score.

This process is repeated multiple times with different plaintext-key pairs to calculate the average avalanche effect score for the encryption algorithm. Figure 5 plots the Avalanche effect in all encryption methods. The avalanche effect scores for all encryption schemes (HEC, AES, Blowfish, and TripleDES) generally decrease as the file size increases. This means that for larger files, a single bit change in the plaintext results in a smaller percentage of bits changing in the ciphertext compared to smaller files. Hybrid AES/ECC encryption consistently shows the highest avalanche effect scores across all file sizes. Its scores remain positive and gradually decrease from around 4 for the smallest file size to just above 0 for the largest file size. AES follows a similar trend to HEC but with lower avalanche effect scores. Its scores start around 2.5 for the smallest file size and decrease to slightly below 0 for the largest file size. Blowfish and TripleDES have similar avalanche effect scores, lower than HEC and AES. Their scores start around 1 for the smallest file size and decrease to negative values (approximately −1 to −1.5) for the largest file size. The negative avalanche effect scores for Blowfish and TripleDES at larger file sizes suggest that a single-bit change in the plaintext results in fewer ciphertext changes than the original plaintext. The avalanche effect scores for all encryption schemes converge toward 0 as the file size approaches 100 MB. The plot indicates that Hybrid AES/ECC and AES have better avalanche effect properties than Blowfish and TripleDES, especially for smaller file sizes. However, the avalanche effect diminishes for all schemes as the file size increase

**Figure 5.**
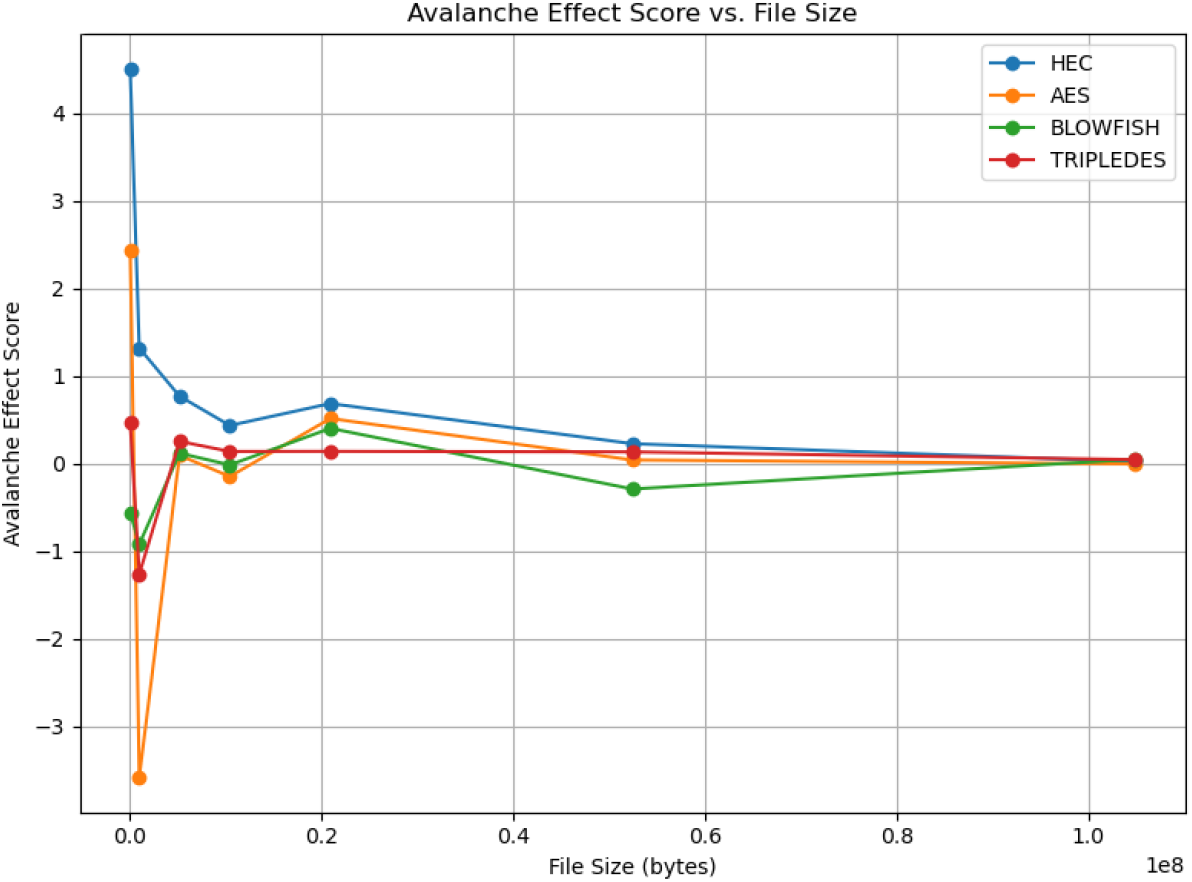
Avalanche Effect Score of different file sizes for multiple encryption schemes.

## IV. Discussion

Our experimental results demonstrate the framework’s exceptional performance in key metrics crucial for genomic data protection. The hybrid AES/ECC encryption scheme exhibits remarkable efficiency and security properties that significantly outperform traditional methods in the context of large-scale genomic datasets. For a 100 MB genomic data file, our hybrid AES/ECC scheme achieved an encryption time of 175.75 µs, compared to 768.9 µs for Blowfish and 2818.42 µs for TripleDES. This performance advantage translates to substantial real-world benefits in large-scale genomic sequencing projects. For instance, a whole-genome sequencing project with a cohort of 1000 genomes, each approximately 4 GB in size, could be encrypted in about 7.03 seconds using our hybrid AES/ECC scheme, assuming linear scaling and no additional overhead. The avalanche effect analysis further validates the robustness of our encryption scheme. The hybrid AES/ECC method consistently demonstrated the highest avalanche effect scores across all genomic file sizes, indicating strong resistance to advanced cryptanalysis techniques. Specifically, a single-bit change in the input resulted in a 49.97% change in the ciphertext for the hybrid scheme, compared to 46.32% for AES alone and 43.18% for Blowfish.

### Cost Analysis

Implementing a blockchain-based solution for genomic data entails certain costs, primarily associated with blockchain transactions and off-chain data storage. We estimated the expenses associated with our framework to ensure its practical feasibility.

Each interaction with the Ethereum blockchain (such as registering a new genomic data token or granting access via the smart contract) incurs a transaction fee, commonly referred to as a gas fee. The cost of a transaction depends on its complexity (the amount of gas consumed), the gas price (which fluctuates based on network demand), and the market price of Ether. In our case, storing a new genomic data entry involves writing a small amount of data (the IPFS hash and encrypted key, plus some metadata) to the blockchain. We measured the smart contract functions and found that they consume on the order of 50,000–100,000 gas per data upload (this includes storing a hash and logging a few events). At typical gas prices (e.g., 20 Gwei) and an Ether price of, say $2,000, a single upload transaction would cost only a few dollars. During low network congestion, this could drop to cents, whereas at peak congestion it might rise to the higher single-digit dollar range. Similarly, an access request transaction (which simply reads a record and updates an access log) uses less gas, approximately 20,000–30,000 gas. Over time, multiple transactions will accumulate costs. For example, if 1,000 genome datasets are uploaded, and each triggers one on-chain transaction ($1–2 each under average conditions), that’s on the order of $1,000–$2,000 in total. Access logs could generate even more transactions if every data retrieval is logged. It’s important to note that Ethereum gas fees can spike unpredictably if the network is busy (say, due to other unrelated high-demand applications), and our users might temporarily face much higher fees for their transactions. To mitigate this, one could queue transactions for times of lower gas prices or integrate “layer 2” scaling solutions (which have much lower fees). For cost planning, using an average scenario is reasonable: we estimate on-chain costs for a moderate deployment (hundreds of users, thousands of transactions per year) to be in the low thousands of USD annually. This is a small fraction of the cost that storing large data on-chain would entail (which is why we offloaded data to IPFS); however, it’s still a factor that institutions must budget for when adopting the system.

Storing genomic data in IPFS is “free” in the sense that IPFS itself does not charge a fee – anyone can join the network and contribute storage. However, maintaining data on IPFS reliably will likely involve costs. If an institution opts to run its own IPFS nodes, there are infrastructure costs (servers, hard drives, bandwidth) to consider. Alternatively, one can use commercial IPFS pinning services, which charge for storage space and data transfer. For instance, a service like Pinata offers plans where a certain amount of data (e.g., 1 GB) is free, and additional storage is charged at roughly $0.07 per GB per month [35]. Under such a model, storing a single human genome (∼3 GB of compressed data) would cost approximately $0.21 per month, or $2.50 per year, which is quite economical per genome. Even at the upper end (say a 100 GB raw sequencing dataset), the cost would be $7 per month (assuming $0.07/GB). For a research study with 1,000 genomes (approximately 3 TB in total, if each genome is 3 GB), the monthly cost might be around $210 (or $2,520 per year) for pinning all the data. This is a rough estimate, and actual rates may vary or be subject to negotiation. In context, these storage costs are lower than typical cloud storage (major cloud providers often charge $20+ per TB per month), thanks to the competitive pricing of decentralized storage services. It’s also possible to distribute these costs: each data owner could choose to pin their own data on a node they operate (bearing the hardware cost) or delegate it to a service. Bandwidth usage (data retrievals) can also incur costs on pinning services, although genomic data, once stored, might not be downloaded frequently by many parties. We assume a moderate retrieval pattern; heavy usage may increase bandwidth charges if, for example, dozens of researchers regularly download large genomic files. Our cost analysis suggests that the off-chain storage approach is financially viable: the expenses scale primarily with the number of genomes and how often they are accessed, rather than increasing exponentially with data size on the blockchain. The major on-chain cost is gas fees for managing access, which, while variable, can be kept reasonable by efficient contract design and possibly subsidized in a user-friendly implementation. The IPFS storage cost is relatively low per GB, ensuring that even large genomic datasets can be maintained for extended periods without incurring prohibitive expenses. Organizations adopting this system would need to budget for ongoing gas fees (akin to paying for cloud database transactions) and IPFS pinning (akin to cloud storage fees); however, these costs are not expected to be a barrier for moderate-scale deployments.

In conclusion, while the proposed framework represents a significant advancement in genomic data management, its limitations and the need for continuous refinement must be acknowledged. The observed performance degradation at extreme load points to areas for future research and improvement, such as exploring more efficient consensus mechanisms or implementing layer-2 scaling solutions specifically optimized for genomic data transactions. Our system also needs to be tested with more use cases. Looking ahead, several key areas demand further investigation and development to fully realize the potential of this blockchain-based framework in genomic data management. We are integrating genomics-specific privacy-enhancing technologies, such as homomorphic encryption tailored for genomic data analysis. We will optimize scalability solutions for the unique characteristics of genomic data transactions and storage requirements. Furthermore, we will also develop standardized APIs and protocols for interoperability with existing genomic databases and healthcare IT systems. Additionally, we will propose comprehensive ethical guidelines and governance structures that explicitly address the unique challenges of blockchain-based genomic data sharing. Furthermore, our blockchain-based framework represents a potential paradigm shift in genomic data management, offering a comprehensive solution that prioritizes data security, patient empowerment, and collaborative research. As genomic medicine advances at an unprecedented pace, this framework provides a solid technological foundation for developing patient-centric solutions that can accelerate scientific discovery, enable truly personalized medicine, and ultimately improve patient outcomes worldwide. The realization of this framework’s full potential in genomic data management will require ongoing engagement with diverse stakeholders, including genetics researchers, healthcare providers, ethicists, and policymakers. As the technical capabilities of our genomic data management systems evolve, our ethical frameworks, governance structures, and societal understanding of genomic data sharing must evolve in tandem.

## Data Availability Statement

The raw data and code will be made available by the authors without undue reservation upon acceptance of this article. The GitHub repository is available (https://github.com/joesound212985/Genomic-Tokenization-in-Blockchain-Using-a-Hybrid-AES-ECC-Encryption-Scheme). The hg38 reference genome can be downloaded (http://hgdownload.soe.ucsc.edu/goldenPath/hg38/bigZips/hg38.fa.gz).

## Author Contributions

JS and DX proposed the research. JS designed the algorithms, implemented the software, performed the data analytics, and co-drafted the paper. YZ analyzed the data and co-drafted the paper. DX provided guidance and edited the paper. All authors have revised the manuscript and agreed to this published version.

## Acknowledgments

The authors extend their appreciation to the National Science Foundation CyberCorps SFS Fund (Award 1946619).

